# Local network determinants of spontaneously emerging cortical maps

**DOI:** 10.1101/764068

**Authors:** Tomer Fekete, David B. Omer, Amiram Grinvald, Cees van Leeuwen

## Abstract

Spontaneously emerging cortical maps related to the functional architecture of visual cortex have been observed initially in anesthetized cats and, subsequently, in monkey, albeit only under certain anesthetic regimes, and not in the awake state. Here we propose a network model that can accommodate these diverse findings. The model identifies two crucial determinants for the emergence of spontaneous map-like activity – local balance between excitatory and inhibitory activity, and the strength of feature-specific synaptic connections (e.g. orientation, ocularity). Our model further shows that dynamically, map-like activity patterns could be triggered either by standing or travelling waves, a mode of operation which is determined by the spatial extent of lateral connections within a given network. Our results suggest that careful pharmacological intervention can unveil the prevalence of maps – recurring spatial patterns of inhomogeneous lateral connectivity - in cortex without the need to explicitly identify area specific optimal features.

## INTRODUCTION

In a series of pioneering studies using voltage sensitive dye imaging (VSDI) in cat primary visual cortex, spontaneous activity under anesthesia was shown to exhibit rich spatiotemporal dynamics. In absence of external stimulation, activity patterns resembling evoked orientation maps often emerged ^1-4^. The same was subsequently demonstrated in monkey, where - not only orientation maps, but also ocular dominance maps arose spontaneously^5^.

These results might suggest that the cortex utilizes a restricted alphabet of “states” (activity patterns) - akin perhaps to symbols - that could be recruited on the fly to meet computational demands ^6^. Arguably, however, under anesthesia the cortex is not engaging in any computational task whatsoever. It is therefore hard to see why cortical networks would engage in computationally meaningful activity (namely expressing bone fide cognitive symbols) that serves no utility. Rather, under anesthesia cells undergo stereotypical changes in membrane potential ^7,8^. Perhaps, therefore, anesthesia coaxes brain networks into simple stereotypical behavior, a behavior of which spontaneously emerging maps are simply a byproduct, rather than indicative of cognitive mechanisms per se. Indeed, under anesthesia long term communication appears to be compromised, and hence the possibility of integration of information is as well ^9^.

We therefore consider whether spontaneously emerging map-like activity patterns could be explained by the specific conditions induced by certain anesthetic protocols – e.g. changing the balance between excitation and inhibition. To this end we compared the data collected in the original primate study by Omer et al. 2018^5^ to equivalent spontaneous voltage sensitive dye imaging (VSDI) data collected under a different anesthetic protocol. In addition, we introduce a detailed biophysical model, which reproduces salient features of the dynamics observed in the original study ^5^. Our model enabled us to examine the local network determinants at the root of the emergence of these activity patterns.

## RESULTS

Our goal in this study was to model the emergence of spontaneous cortical maps in early sensory cortices, taking a highly integrative approach: unlike previous models that focused mainly on reproducing the frequency distribution of emerging map-like activity patterns c.f. ^10^, we wanted to do so while remaining faithful to the actual circuit dynamics. To this end we employed a more biophysically realistic model, our previously introduced generalized Jansen-Rit model (JRM_gen_; see table 1 for model parameters and Methods for a detailed description)^11^. The JRM_gen_ comprises a set of interacting mini-column like components (Figure 1). Each mini column consists of three elements, representing three neuronal pools - pyramidal cells, local inhibitory neurons and local excitatory neurons – which interact locally. The pyramidal cells in turn project to other pyramidal neurons with spatially limited “lateral” connections, relayed through the local excitatory neurons.

**Table 1:**
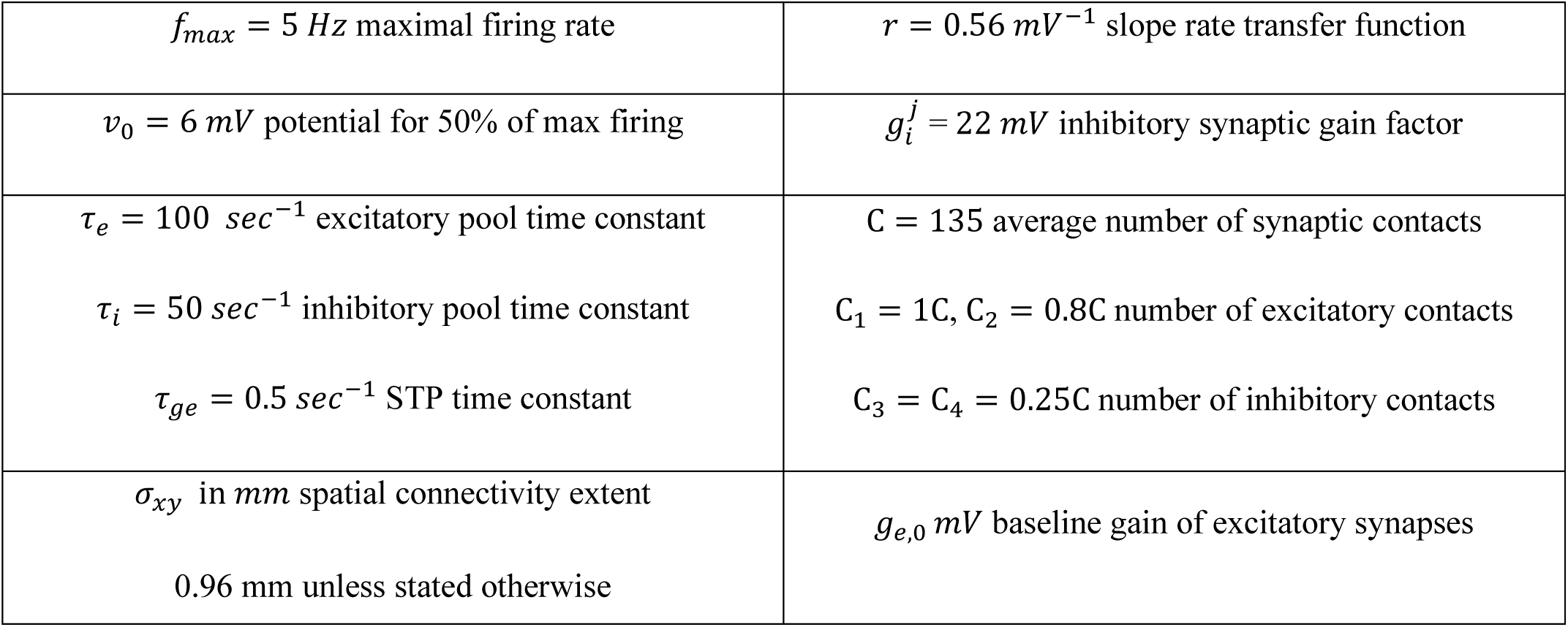
parameters of JR modules.

**Figure 1:**
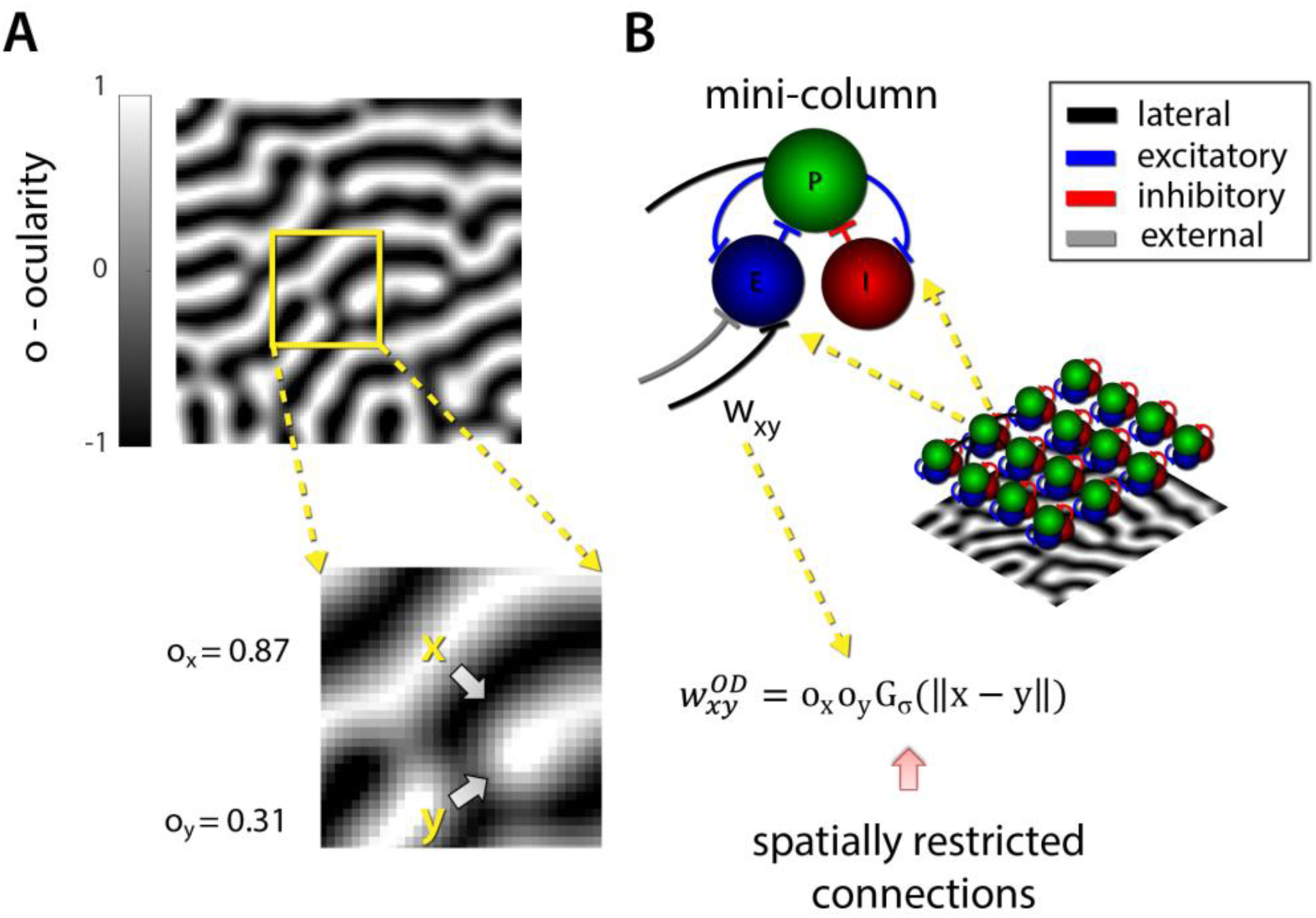
A schematic representation of the V_1_ JRM_gen_. (A) modeling ocularity specific lateral connectivity. An emperical VSDI derived OD map was rectified to be equally distributed between −1 and 1 (representing selectivity for the L and R eye). Ox an Oy are the ocularity preference values of the loci marked x and y on the cortical surface. (B) The V_1_ JRM_gen_ comprises an array of mini-colums following the schematic diagram above. Each mini-column is coupled to other mini-colums in a spatially restricted manner (governed by σ_xy_) by w_xy_. The OD contribution to coupling between locus x and y on the cortical sheet is given by 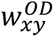.

To adapt this generic model of a cortical area to represent the circuitry of primary visual cortex, we followed the approach introduced in^6^, and used actual imaging data based functional maps to derive the simulated network connectivity patterns. Further still, to try and ensure the generalizability of our results, we chose to also model the recently reported state-dependence of the emergence of spontaneous map-like activity^10^. To fully clarify motivate our modeling choices we will begin by presenting the abovementioned features of spontaneous neural dynamics, as captured by VSDI, that we aimed to model.

We revisited the data reported in Omer et al. 2018 ^5^, in which an anesthetic protocol involving opioid anesthetics (Remifentanil; which we will refer to simply as Remifentanil – see methods for details) led to spontaneously emerging ocular dominance (OD) map-like activity patterns, as well as orientation map-like patterns in the primate (*Macaca fascicularis*). Figure 2 shows an example of a spontaneously arising OD-map like activity patterns. This example suggests that spontaneous OD maps can propagate along the surface of primary visual cortex, and that their emergence can be associated with a “dampened oscillation” pattern - a map like activation pattern followed by its inverse, albeit with reduced expression, in turn succeeded by the original pattern even more diminished.

**Figure 2:**
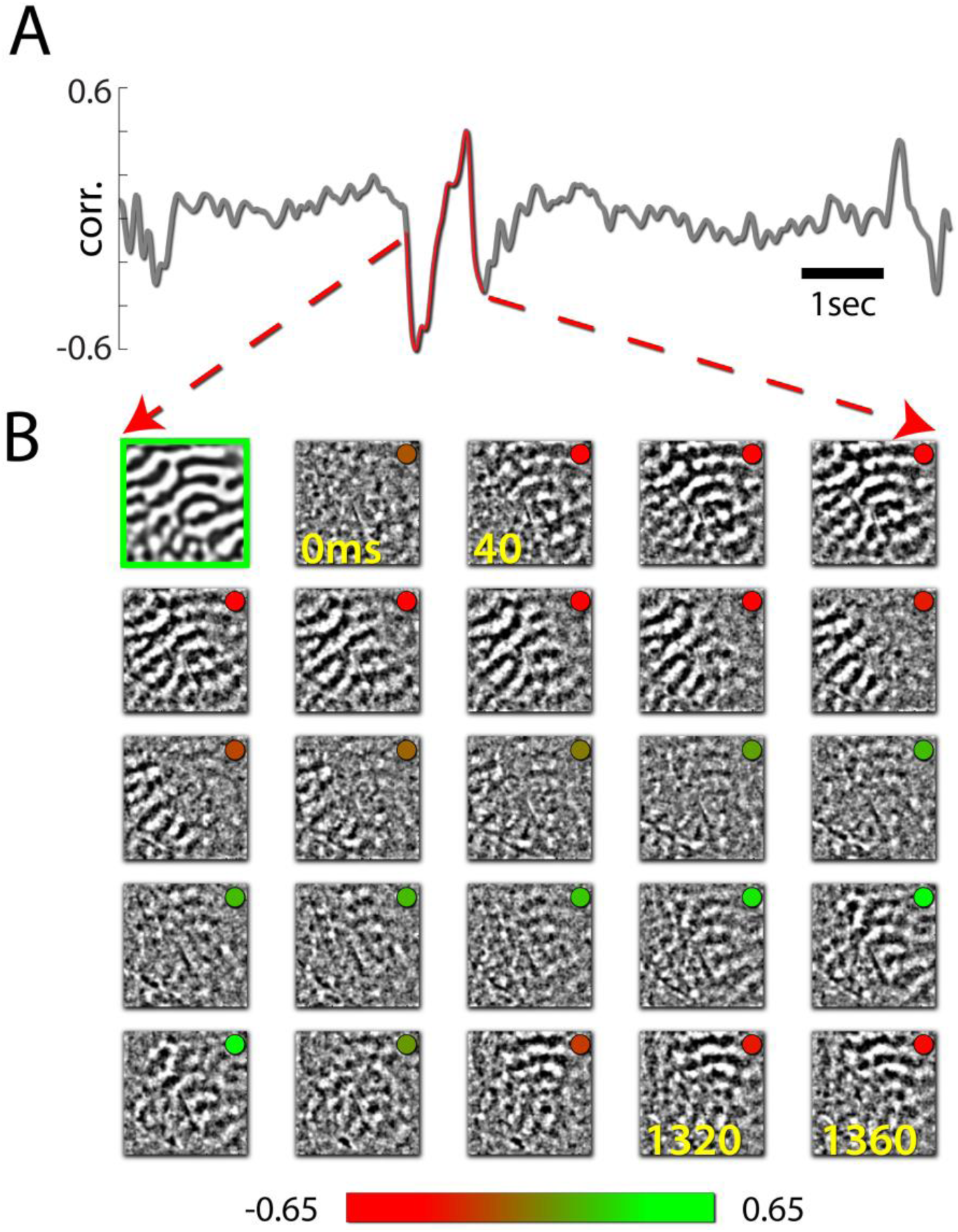
Spontaneous ocular dominance (OD) dynamics under AP2. A) the frame by frame correlation between the OD map and spontaneous activity patterns in primary visual cortex of an nesthetized monkey. B) The high pass component of the optical frames corresponding to A (first frame on the top left is the OD map framed in green for reference, see also fig. 1 in Omer et al. 2018).

Carrying a frame by frame correlation analysis on these data and comparing these ongoing activity patterns to evoked OD and orientation functional maps, we then identified local maxima in the resulting correlation time series, and used those time points as triggers for averaging the optical frames surrounding these events. On average, map-like patterns were enabled by depolarizing events, and were expressed mostly during the high phases of such events (see also Omer et al. 2018)^5^. In this example, on average, spontaneous map-like states appeared to propagate as waves traveling on the anterior posterior cortical axis (Figure 3).

**Figure 3:**
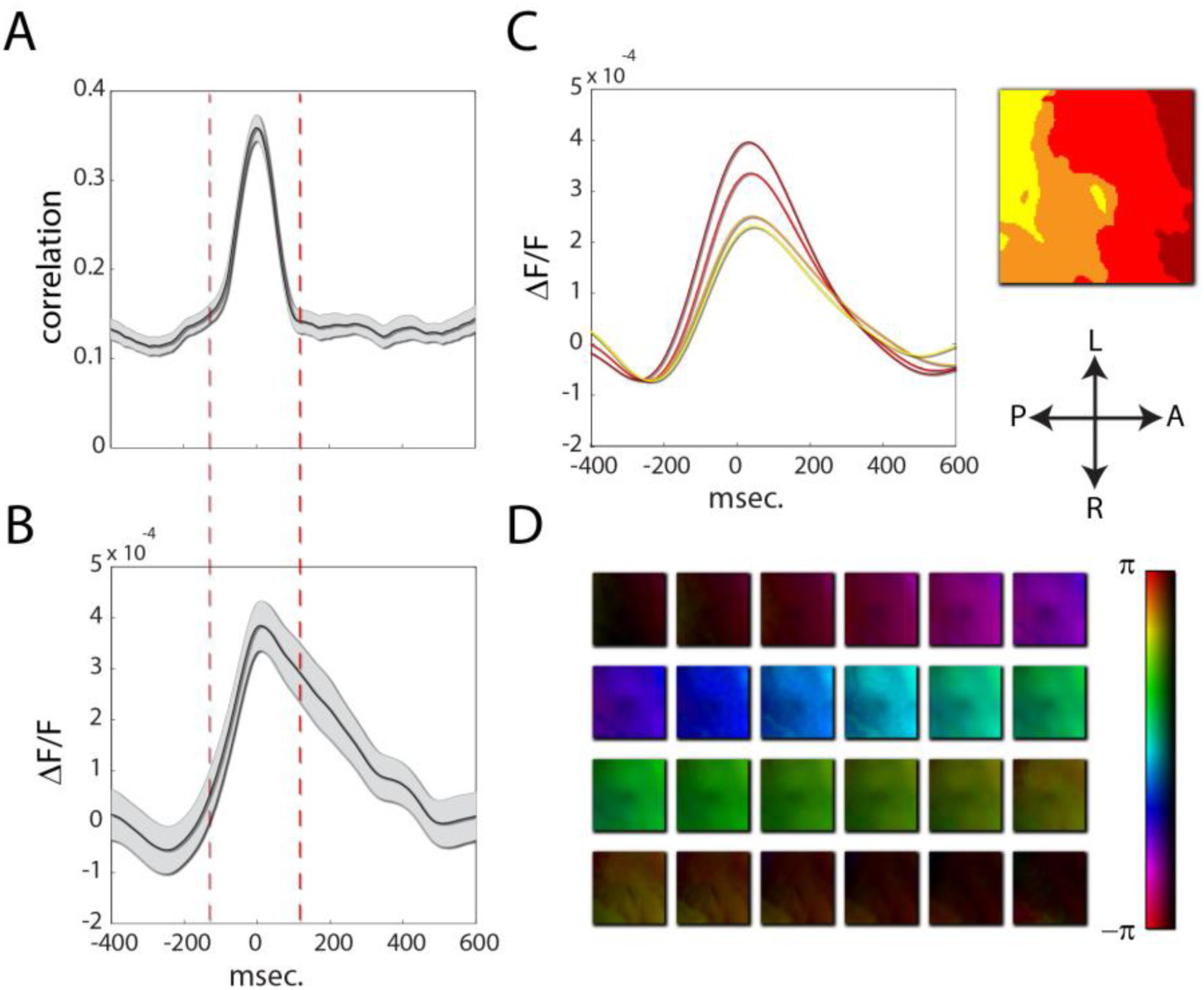
Spontaneous emerging maps typically propagate as waves. Frame by frame correlation to map time series were used as triggers for averaging. A) The average correlation event. The top 5% of local minima in the frame by frame correlation were used as triggers. B) the average amplitude of the VSDI signal across all pixels in the imaged area during spontaneous map emergence. Depolarization preceded map emergence C) the average optical time series (one for each pixel) were clustered according to similarity to produce four clusters. The average time series for each cluster were color coded (left), and the pixels corresponding to each cluster overlaid on the cortical sheet (right). On average, depolarization propagated from the anterior to the posterior regions of visual cortex during map emergence. D) The average optical frames transformed via the Hilbert transform after filtration to the θ+δ band, exposing the wave event associated with map emergence.

In the monkey, OD map like states were almost twice more prevalent than orientation ones - spontaneous activity frames were correlated to the three functional maps derived in each experiment (left and right OD and cardinal and oblique orientation maps) and were classified as one of the four states in a winner take all fashion (figure 4A). Additionally, the average of the frame by frame correlation to the respective map was computed for each class of frames, and OD like states were found to be more pronounced (figure 4B; see also Omer at al. 2018^5^, fig. 3)

**Figure 4:**
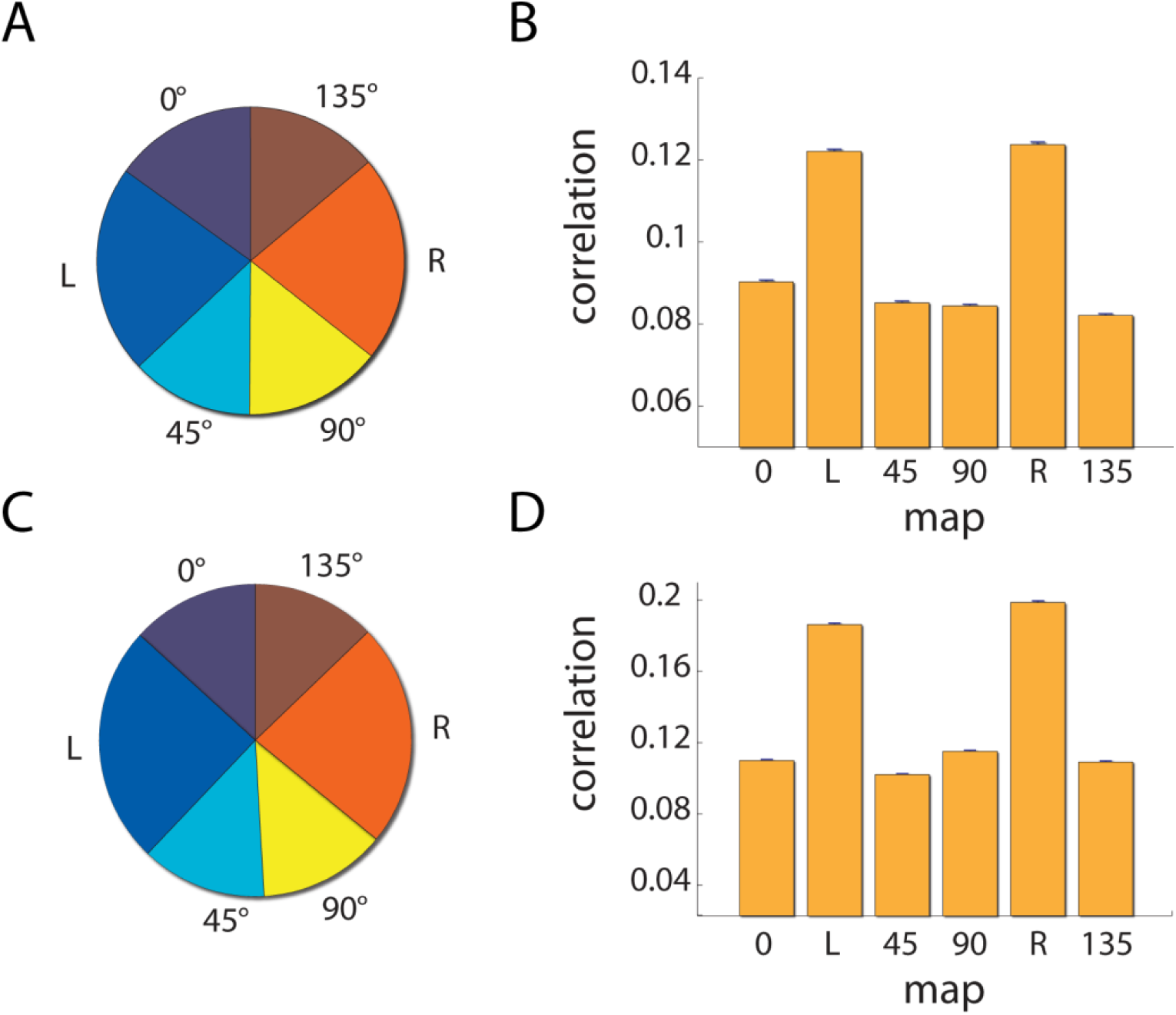
Mixed connectivity neural mass model reproduces frequency of occurrence and salience of spontaneous map-like patterns in monkeys. Top row: empirical data taken from Omer et al. 2018. Bottom: simulated data A) The distribution of spontaneous map-like states. OD states were more prevalent than orientation states. B) The average correlation for map-like spontaneous states. OD-like spontaneous patterns were more strongly correlated to the OD map. C) The distribution of spontaneous map-like states for a simulated cortical sheet in which both orientation and OD connectivity were embedded (see text for details). D) The average correlation for map-like spontaneous states in the simulated data. OD like spontaneous patterns were more strongly correlated to the map, as compared to spontaneous orientation states.

As was observed in anesthetized cat ^4^, activity patterns highly resembling the functional maps emerged as leading principal components (figure 5), and the prevalence of OD-like states was reflected in the increased variance explained by the associated principal component as compared to the components associated with both the cardinal and oblique orientation maps.

**Figure 5:**
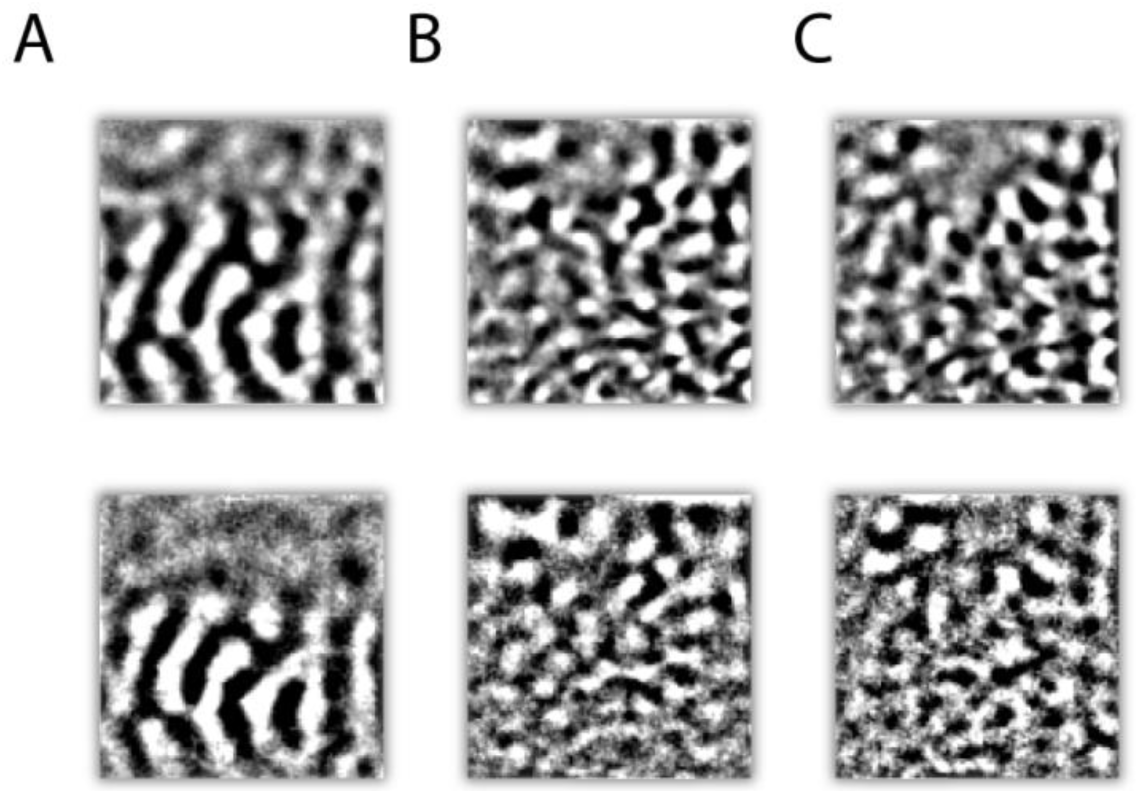
Functional map-like patterns emerge as principal components of spontaneous fluctuations under Remifentanil anesthesia in the primate. Top row: the OD and cardinal and oblique functional maps. Bottom row: leading principal components of data collected from primate primary visual cortex under Remifentanil anesthesia sorted according to their explained variance.

However, as reported by Omer et al 2018^5^, spontaneous map like activity patterns were not observed in quiet wakefulness. One possible explanation for this is that spontaneously emerging map-like activity is triggered by reduced cortical inhibition induced by the anesthetic drug. We therefore analyzed the data of VSDI experiments using a Ketamine based anesthetic protocol (hereafter Ketamine – see methods for details), a substance which in anesthetic doses is known to reduces cortical synchrony ^12^, and hence arguably increases cortical inhibition. Under Ketamine as in the awake state^5^, we did not observe spontaneously emerging maps. As a proxy for determining the level of cortical inhibition in each of the three states, we carried out pixel pair-wise correlation analysis to measure cortical synchrony. As can be seen in Figure 6 synchrony under Remifentanil was increased as compared to quiet wakefulness, whereas Ketamine reduced synchrony.

**Figure 6:**
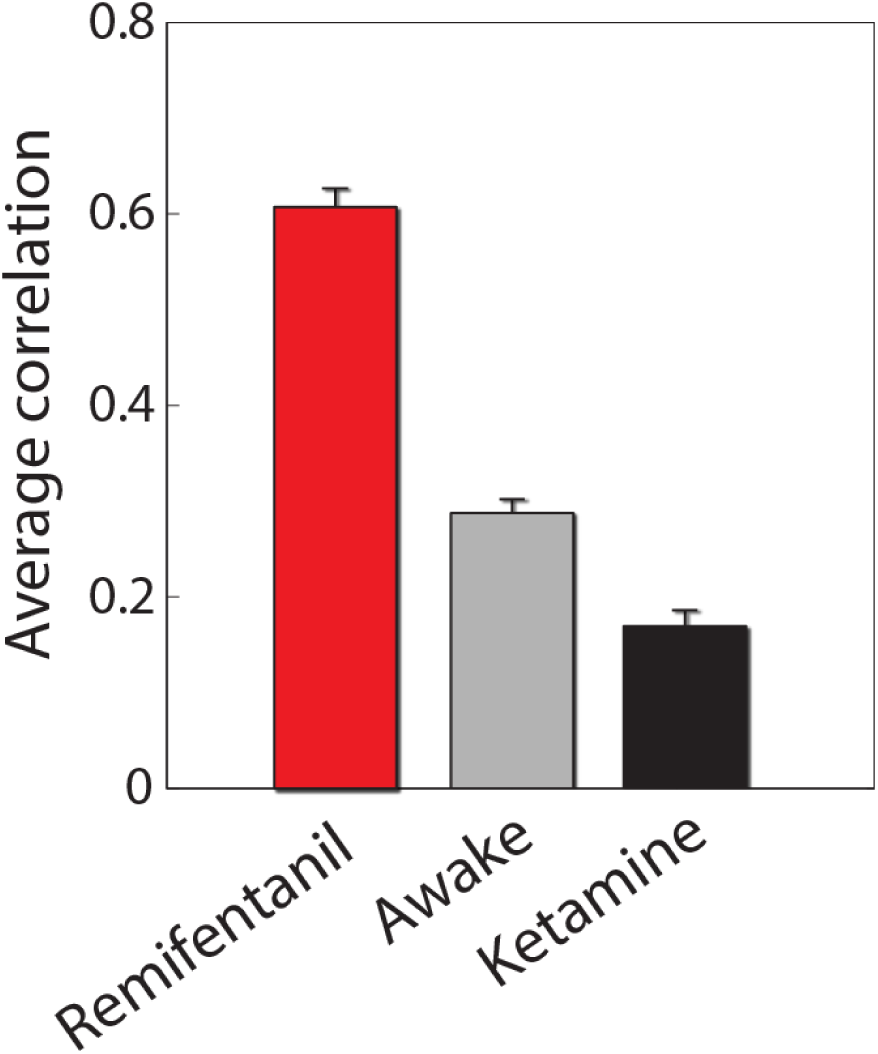
Effect of anesthesia on cortical synchrony. For each spontaneous sequence of optical frames, the average correlation between each pixel, and the pixels in a 10-radius neighborhood was derived and averaged across conditions. While Remifentanil greatly increased synchrony as compared to quiet wakefulness, Ketamine resulted in reduced synchrony, arguably attesting to higher than normal levels of cortical inhibition

We employed a neural mass modeling approach to try and recreate the salient features of the data, simulating spontaneous activity by driving a model of a cortical sheet by independent noise fluctuations. Connectivity in the model reflected both ocular dominance and orientation selectivity in a fashion dictated by empirical functional maps (see methods). We found that to observe spontaneously emerging map like activity patterns, several conditions had to be met (see figure 7): 1) the strength of feature specific lateral connections had to be sufficiently large – but not exceedingly so, otherwise the network enters a bistable regime 2) the balance between excitation and inhibition had to be sufficiently tipped towards excitation (that is, the cortex disinhibited). 3) the spatial extent of lateral connectivity had to be widespread enough (at least ∼ 1mm).

**Figure 7:**
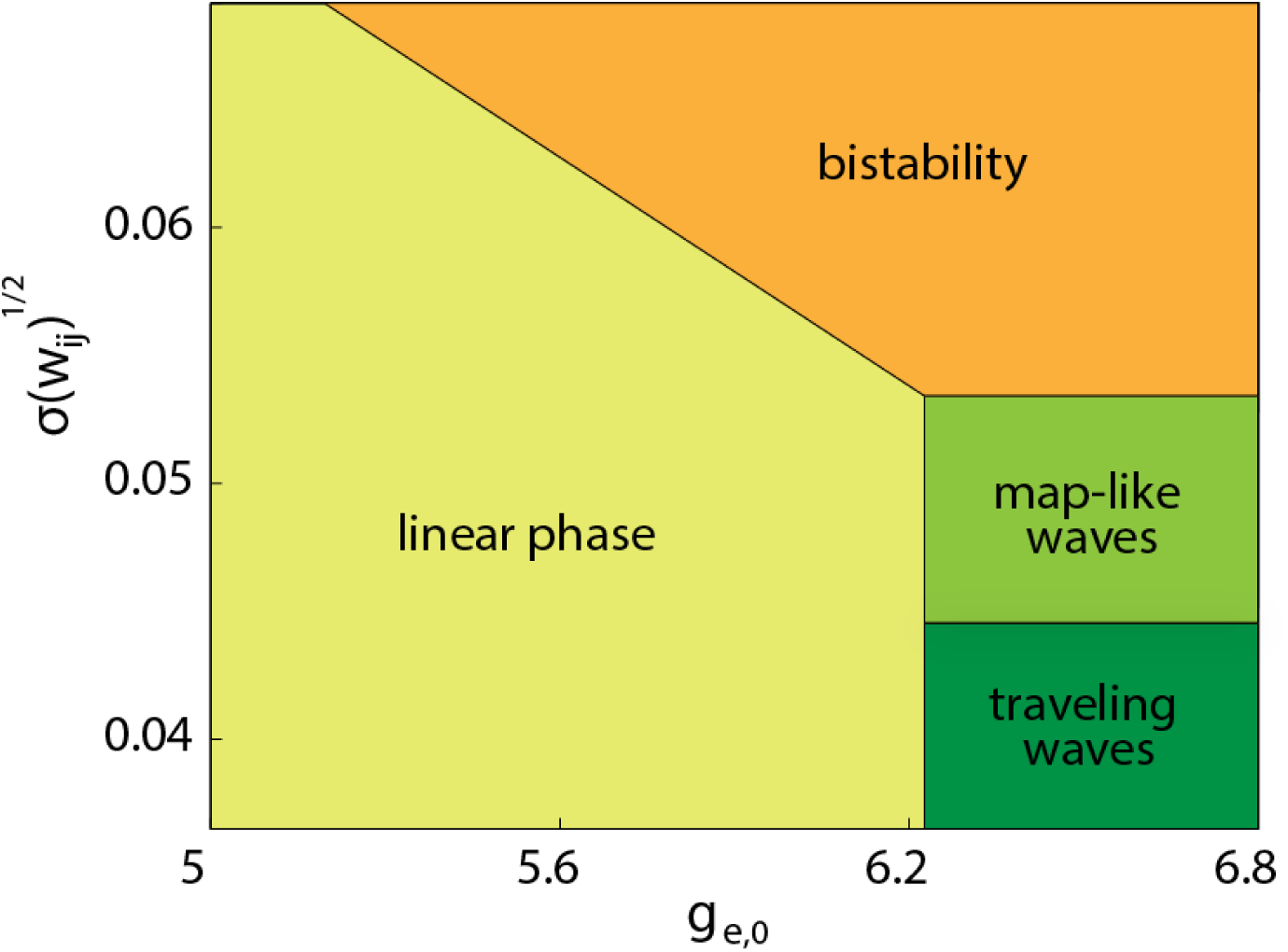
Emergence of map-like activation patterns depends on cortico-cortical inhibition and efficacy of feature selective synapses. Approximate phase diagram for the simulated cortical sheet. Four dynamical regimes were identified for the simulated network, which were triggered by the excitation to inhibition ratio g_e,0_, and the square root of the standard deviation of lateral connection magnitude: A linear phase, in which the network exhibited (colored) noise oscillations. A pure traveling wave phase, induced by reduced local inhibition. A map like activity regime, which requires increased strength of lateral connectivity. A bistable regime, in which excessively strong lateral connections lead the simulated network to alternate between (near) orthogonal map like patterns. Borders between dynamical regions were not sharp, but rather probabilistic.

Figure 8 shows an example of a simulated cortical sheet in which the above conditions were met, leading to the emergence of spontaneous activity maps. In this example a spontaneous OD-like pattern emerges. The simulated emergent map-like activity followed a dynamics similar to that observed in real data (compare to figure 2), propagating along the cortical surface, as a dampened oscillation. As can be seen in figure 8C, the spontaneous map was triggered by a traveling wave. If the spatial extent of lateral connections varied from ∼ 1mm by 0.4mm, either fragmented patterns emerged, or maps were expressed as standing waves (smaller and larger values respectively).

**Figure 8:**
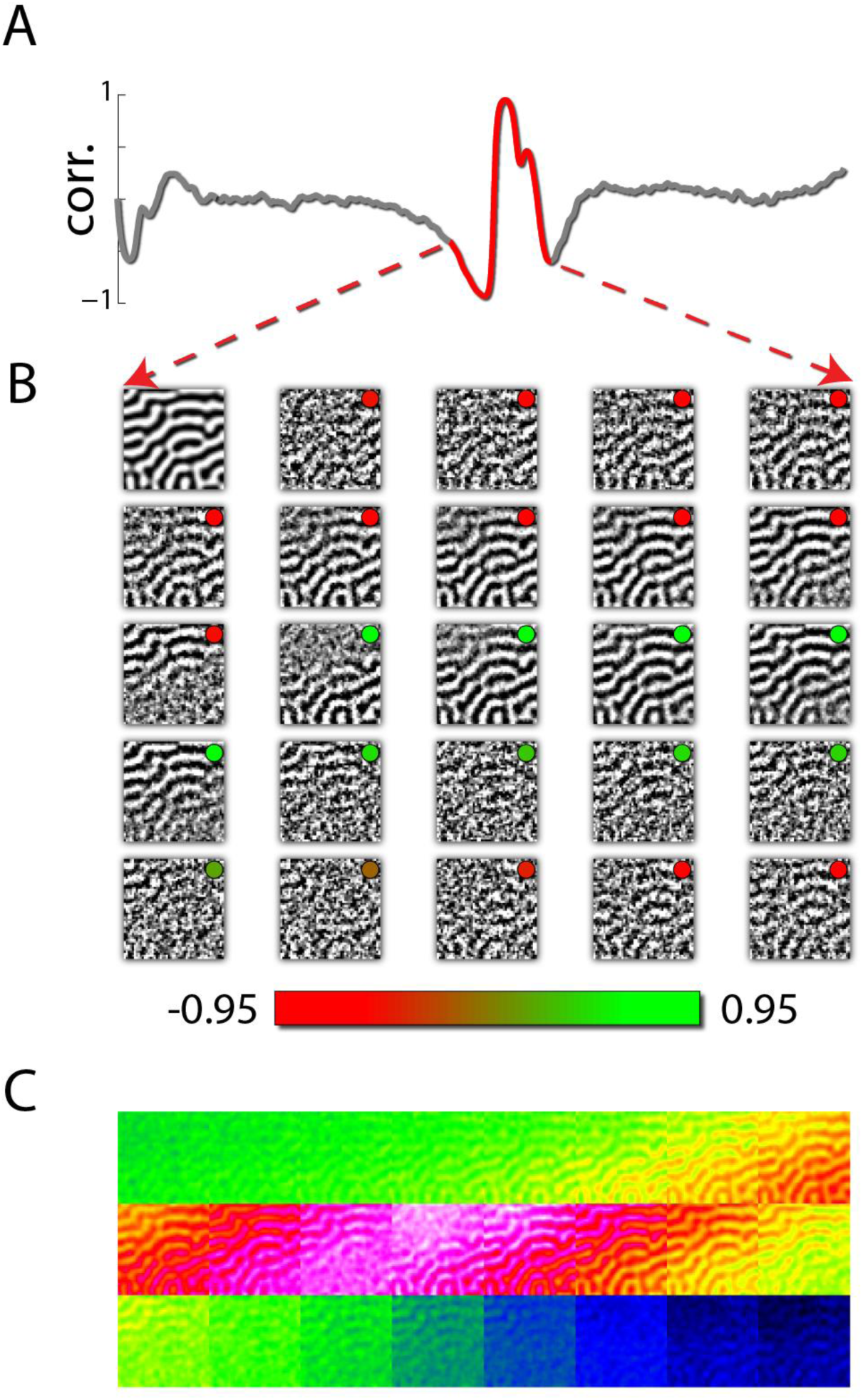
Spontaneous emergence of OD maps in a neural mass model simulation propagate as a traveling wave. A) the frame by frame correlation between the OD map and spontaneous activity patterns in the simulated cortical sheet. B) the corresponding activity patterns. Note the similarity to 1A. B) The (high pass component of the) simulated activity patterns corresponding to A C) the unfiltered frames

In the network shown in figure 8, the ratio between OD and orientation absolute synaptic strength was 0.96, the ratio between their respective standard deviation 0.72, and the average contribution of OD synaptic tracts was 0.0004. This resulted in a distribution of relative frequency of OD and orientation like patterns which was nearly identical to the empirical one (figure 3C). Additionally, the average correlation for each class of patterns followed a similar distribution to the one observed in the actual data (figure 3D).

Finally, we also carried out principal component analysis on these data. As in the empirical data, leading principal components (in the simulated data the first three) highly resembling the functional maps emerged, with the OD map followed by the cardinal and then oblique maps (figure 8 and compare to figure 4). A previous modeling study^13^ showed that in simple rate networks driven by noise, in low noise regimes the principal components of such “spontaneous” fluctuations are expected to be identical to the principal component of the synaptic connectivity matrix (the pattern of lateral connections). As also shown in a previous study^4^, we found that even though our model is much more biophysically realistic (and hence complicated), and even though the noise driving the system was high amplitude, the leading principal components of the connectivity were near identical to those of the spontaneous data (figure 9).

**Figure 9:**
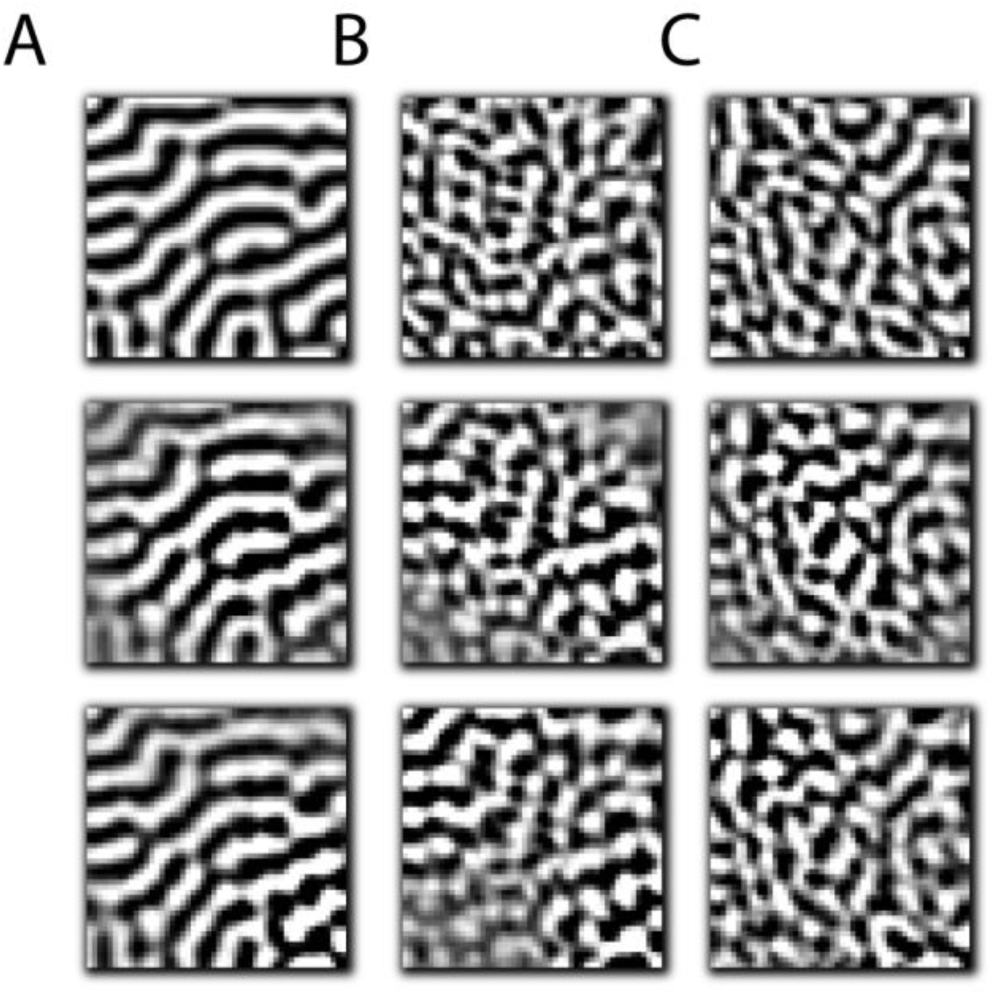
spontaneously emerging maps reflect the principal components of lateral connectivity. Top row: the functional maps used in deriving the connectivity of the simulated cortical sheet (see text for details) Middle row: leading principal components of the connectivity matrix sorted according to explained variance. Bottom row: the leading principal component of spontaneous data simulated through driving the network with noise sorted according to explained variance. As can be seen the principal components of the spontaneous fluctuations and the connectivity matrix are virtually identical, and highly similar to the functional maps (up to edge effects)

## DISCUSSION

In this study we set out to explore the hypothesis that spontaneously emerging functional map-like activity patterns in primary visual cortex, rather than being markers of typical cognitive processing, are in fact byproducts resulting from specific pharmacologically induced changes to the underlying network, such as reduced cortico-cortical inhibition. Indeed, prima facie support for this idea is found in the fact that such patterns were only observed in anesthetically induced high synchrony states, and not in normal and low states of synchrony found in quiet wakefulness and under different anesthetic protocols.

To simulate spontaneous activity in primary visual cortex we employed a biophysically realistic model at the mini-column level. To this end we extended our previously introduced generalized Jansen-Rit model (JRM_gen_)^11^, by incorporating more realistic connectivity, namely both orientation and ocular dominance spatially limited synapses in the lateral connectivity pattern. These connections were derived from actual primate OD and orientation maps^5^. Our V1 specific JRM_gen_ offers significant advantages over previously suggested models for spontaneously emerging maps. One major such advantage is that it produces much more realistic spontaneous dynamics as compared to e.g. ^6,10^ – just like the single JRM it can generate activity which exhibits a 1/f spectrum, produce oscillations in specific bands such as α, and give rise to realistic EEG-like epileptic activity if EIB is increased sufficiently. Further still, it extends the approach of ^6^ to model spatially limited orientation sensitive lateral connections, to OD synapses as well. To model the effect of different anesthetic protocols we varied two parameters – the EIB (controlled by the parameter *g*_*e*,0_ which represents the gain of excitatory synapses), and the strength of feature specific synapses (controlled by the parameter *σ*(*w*_*ij*_) – the standard deviation of synaptic efficacy).

The V1 JRM_gen_ reproduced salient features of spontaneously arising functional-map like activity patterns: map-like patterns were triggered by spontaneous depolarizing events propagating as traveling waves. When synaptic weights were properly adjusted, spontaneous activity exhibited both OD as well as orientation map-like activity patterns, with a similar salience (in terms of correlation to the evoked maps) and frequency to that observed empirically. Finally, as was the case for the empirical data, the leading principal components of spontaneous fluctuations highly resembled the functional maps, with the OD map leading first the cardinal and then the oblique orientation maps - as can be expected from the relative frequency of their spontaneous expression.

Spontaneous map-like activity patterns emerged only if the balance between excitation and inhibition was sufficiently tipped towards excitation. In other words, our model supports the idea that cortico-cortical disinhibition is at the root of this intriguing phenomenon. This result strongly suggests that spontaneous map-like activity is not part of the cortical strategy for meeting computational demands - e.g. coding for expectations or context – given that a hallmark of cortical activity in the awake state in balance between the two.

An ever-growing body of work underscores the crucial role that spontaneous brain activity plays in action, perception and cognition ^1,14-38^. At first blush it might seem that the results we report here indicate otherwise. However, the exact opposite is true: if indeed the spatiotemporal structure of spontaneous activity carries information that serves, supports and shapes cognitive processing in accordance with task and situational demands, such structure should desist, when cognitive and perceptual processing are arrested by anesthetic agents. And indeed, our results indicate that the spatiotemporal activity structures produced by very specific anesthesia induced conditions are in a deep sense “trivial”: they are entirely stochastic in igniting, and their temporal structure is stereotypical, and hence they cannot be said to encode the likes of context or expectation.

It is interesting to note the species-specific difference between spontaneously emerging maps in cats ^4^ and monkeys^5^ – in the former case orientation maps were dominant whereas in the latter scenario ocular dominance maps were more prominent. This likely reflects the respective importance that stereoscopic processing plays in each species. Surprisingly, in our model the dominance of OD patterns over orientation ones was not a result of the strength of OD synapses relative to orientation ones: in fact to reproduce the empirical results orientation synapses had to be slightly stronger – OD synapse strength was set to 96% relative to the orientation ones. The OD dominance can be explained by the respective distributions of feature selectivity in primate visual cortex – whereas OD preference is near bimodal, orientation preference in nearly uniform. Therefore, the driving force for OD states is stronger as it is reinforced by larger coalitions, all other things being equal.

An intriguing possibility suggested by our results is to chart cortex for the existence of previously unknown functional maps, in the event that those maps emerge from typical patterns of spatially inhomogeneous lateral connectivity, without the recourse to explicitly determine the optimal features such maps are attuned to. Our model predicts that this can be achieved by inducing sufficiently strong slow wave activity through pharmacologically induced disinhibition. Likely, this can be facilitated by better understanding of the effects compounds have jointly on the effective connectivity of lateral synapses.

To conclude, this study highlights the fact that currently VSDI offers an unrivaled means of exploring the columnar architecture of cortex. The present results call for employing additional anesthetic regimes, as well as specific pharmacological interventions down to the level of transmitter and receptor, which should provide indispensable information about the local structure and function of cortical networks,

## METHODS

All surgical and experimental procedures were approved by the Institutional Animal Care and Use Committee at the Weizmann Institute of Science.

### VSD imaging

The data from the remifentanil and awake experiments that were analyzed here were previously published by Omer et al. 2018^5^. For more detailed information regarding the experimental procedures see Omer et al. 2018 as well as Omer et al. 2013^39^. Long-term VSD imaging was carried out in two adult male Macaca fascicularis monkeys. A head-holder and two cranial windows (25 mm internal diameter) were placed over the primary visual cortex and cemented to the cranium with dental acrylic cement. Several months afterwards, the monkeys underwent craniotomy and the dura mater inside the chamber was resected to expose the visual cortex. The anterior border of the exposed cortex was 3–6 mm anterior to the lunate sulcus. The center of the chamber was 0°–4° below the representation of the vertical meridian in V1 and 2°– 4° lateral to the horizontal meridian. A thin, transparent silicone artificial dura was implanted over the exposed cortex ^40^. The surgical procedure has been reported in detail previously ^41,42^.

At the beginning of each VSDI session the cortex in the imaging chamber was stained with an oxonol VSD, RH-1916. To ensure that the dye solution (0.2–0.3 mg/ml) was sterile, it was filtered through a 0.2-µm filter. After 2 h of staining, the chamber was reopened, and the cortical surface was washed with ACSF until the solution was as clear as the ACSF. At the end of the staining the artificial dura was placed back over the cortex. Hard agar solution was gently poured onto the real dura in the periphery of the cranial window, and a more dilute and transparent agar solution was used to fill the chamber, above the artificial dura. In the awake experiments, during the entire painless staining procedure and preparation time for imaging, the monkeys were awake and sat calmly in their chairs.

For real-time optical imaging, a custom-written program was used to control a MICAM Ultima high-speed camera (SciMedia, Japan), with a resolution of 100×100 pixels. Each pixel viewed 120×120 microns of cortex. The exposed cortex was illuminated using epi-illumination with an appropriate excitation filter (peak transmission 630 nm, width at half height 30 nm) and a dichroic mirror (650 DRLP), both from Omega Optical (Bratlleboro, VT, USA). To collect the fluorescence and reject stray excitation light, a barrier postfilter (RG 665; Schott, Mainz, Germany) was placed above the dichroic mirror. Data were recorded at 200 Hz. To avoid irreversible photodynamic damage to the cortex, each measurement was conservatively terminated after a total of ∼ 5 min of net accumulated illumination time.

To map the functional architecture of orientation columns we used four stimuli, each lasting 50–100 milliseconds, with a minimum inter-stimulus interval of ∼ 150–250 (for detailed explanation of the experimental protocol see Omer et al 2013)^39^. Stimuli were alternated for a continuous period of ∼ 10 s. We repeated the 10-s stimulation session eight times with an inter-session interval of 2.5 s. The visual stimulation consisted of full-field gratings alternated with an inter-stimulus interval (gray screen). The gratings were shown in one of four orientations, and the orientation angle was changed between presentations either in a fixed order (0°→45°→ 90°→135°→0) or randomly. To stimulate the entire imaged cortical area, the spatial phase of each orientation stimulus was shifted between presentations. The screen background was kept isoluminant for the entire trial period, including the period of fixation prior to stimulus onset.

To map the functional architecture of ocular dominance columns we used either full-field isoconcentric rings of flickering checkers or a drifting grating (contrast, 90%; size, 13×13°; spatial frequency, 2 cycles/degree; temporal frequency, 2 degrees/s; mean screen luminance 23 cd/m^2^). A stimulus was presented to one eye at a time, alternating between eyes every 200 milliseconds by the use of a pair of ferroelectric shutters (DisplayTech Inc.). Stimuli were presented on a 21-inch CRT monitor (Sony GDM-F500) at 120 Hz, placed 100 cm from the monkey’s eyes. We combined our custom-made imaging software with the VCORTEX software package (VCORTEX 2.0, NIMH) for controlling behavioral events. The precise time course of the stimulus was monitored by a photodiode and saved as an analog channel that was synchronized to the data acquisition.

For the ongoing activity sessions in the awake state the monkeys were trained to sit quietly for periods of ∼ 11 seconds while their eyes were covered with eye shutters to ensure an even illumination pattern. At the end of each ongoing session the monkey was rewarded with 0.2-0.4 ml of water or juice. In each experiment day a total of ∼ 10-12 trials of 10 seconds each was collected with an inter trial period of ∼ 2-4 seconds. Overall the imaging session was very brief and animal state was constantly monitored with local-field potentials.

Ongoing activity was also imaged in a similar fashion to the awake experiments under two different types of anesthesia: The animals were first anesthetized with a mixture of ketamine (10 mg/kg) and Diazepam (1 mg/kg). Anesthesia was maintained with remifentanil (6-120 microgram/kg/h), and the animal was paralysed using Pancuronium-bromide (Pavulon, 2 mg kg-1 hour-1)., while the monkeys were artificially respirated. In the second anesthetic protocol, anesthesia was maintained with intramuscular ketamine administered as necessary to achieve cortical deactivation (as observed through local field potentials - LFPs). The state of the animals and the level of anesthesia were monitored continuously by measurement of EEG, LFP, electrocardiogram, endtidal CO2, and rectal temperature.

### Data preprocessing

Bleaching artifacts were removed by fitting data with a sum of two exponentials. Next, heart pulsations were removed by removing the ECG triggered average of each time series. Breathing pulsations were removed following the method described in ^43^. Data were temporally smoothed to the 0.2-8Hz band, and spatially filtered with a 600μm HWHH high pass filter. Next, principal component analysis (PCA) was carried out separately for each experimental condition. Due to the vast number of frames PCA was carried out sequentially, similar to batch ICA analyses ^44^: PCA was carried out to each consecutive 1024 optic frames. The 100 leading components from each batch were concatenated, and finally PCA was carried out again, resulting in a basis which we refer to as *signal*. The same procedure was applied to the difference between the raw data and the possessed data resulting in an orthonormal basis we refer to as *noise*.

The resulting bases was used for denoising the data: data ware reconstructed using the *signal* basis and, in order to reduce shot noise, only the components explaining 80% of the variance were used. Rather than using the standard first frame division method *signal* components were projected out of the data in a semi-automatic procedure informed by *noise* components: The correlation coefficient between the *signal* and *noise* components was computed, and *signal* components which were similar to *noise* component (*r* > 0.45) were discarded after visual inspection to verify that they did not contain functional patterns.

### Derivation of functional maps

Maps were computed for each trial using a general linear model: box car regressors were convolved with a custom-tailored impulse response constructed via the SPM8 software (http://www.fil.ion.ucl.ac.uk/spm/software/spm8/). The resulting coefficient maps were then averaged across trials. Next, orthogonal orientation maps/different eye stimulus presentations were subtracted to obtain differential maps (orientation, ocular dominance (OD) maps respectively).

### Correlation analysis

Optical frames collected in the absence of stimulation (ongoing activity) were compared to the evoked maps using the correlation coefficient as a measure of similarity each frame at a time. To enhance the contribution of functional signals to the resulting scores images were smoothed with a Gaussian isotropic kernel of HWHH of 60 μm^2^.

### Simulating spontaneous activity

To model the spontaneous activity of a patch of primate primary visual cortex we employed the generalized Jansen-Rit model (JRM_gen_). The modifications we made to the original model were reported in ^11^ and therefore will only be presented briefly below:

The simulated cortical sheet comprised column like nodes, each following the Jansen-Rit equations ^45^. The JRM is a neural mass model that recreates many observed properties of the EEG signal. It comprises three pools of neurons: pyramidal cells, local excitatory neurons and local inhibitory neurons.

The input to a pool *I*(*t*) is propagated along the dendrites to the cell bodies impacting the membrane potential at the cell body *ν*(*t*) through *ν*(*t*) = *s* * *I*(*t*) where *s*, the “synaptic operator” is given by:

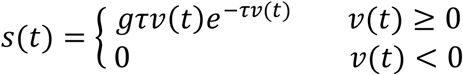

*g* determines the maximal amplitude of post synaptic potentials and *τ* is the membrane time constant.

The corresponding differential equation is

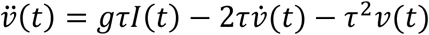

which can be written as a system of two first order equations:

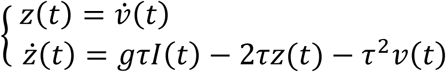

The membrane potential of the *j*^*th*^ pool of pyramidal neurons is denoted by 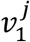, and 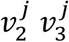 denote the respective quantities for the *j*^*th*^ local excitatory and inhibitory pools.

The pool of pyramidal neurons is driven (regulated) by both local pools of neurons:

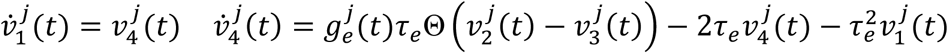

where, as stated above 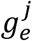 is the synaptic gain factor of the *j^th^* excitatory pool and Θ is the membrane potential to firing rate transfer function

Local excitatory neurons receive pyramidal input as well as input from other sources:

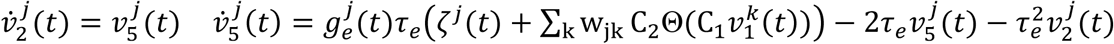

where w_jk_ are the lateral connections between columns, andζ^*j*^(*t*) reflects the firing rates originating from other areas.

Finally, local inhibitory neurons are driven by pyramidal firing:

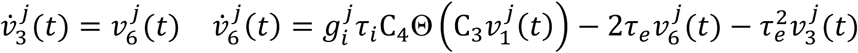

where the constants C_i,_ i = 1 … 4 are proportional to the average number of synapses between populations.

The potential to firing transfer function is given by: 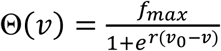, where *f*_*max*_ is the maximal firing rate, *ν*_0_ is the potential for which 50% of the maximal firing is achieved and *r* is the slope of the sigmoid at *ν*_0_.

The output variables of interest are 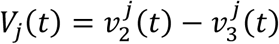 - the membrane potentials of the main family of neurons. We employed the well-understood standard parameters for each module (see table 1) ^46^.

The balance between excitation and inhibition (E/I ratio) is governed by the ratio between 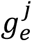 and 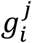. For the sake of simplicity, changing the balance can be effectuated by either increasing or decreasing 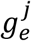 (disinhibition vs. inhibition). In the single column case, sufficiently large disinhibition leads to “epileptic” spiking. However, we have found that in the multi-column scenario it is necessary to turn 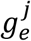 into a dynamic variable that introduces short term plasticity (STP) of excitatory synapses in order to preclude the network from saturating, i.e.:

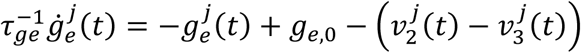

where *g*_*e*,0_ is the baseline excitatory synaptic gain factor, which in effect determines the excitation to inhibition balance, and *τ*_*ge*_ is the time constant determining the temporal scale in which STP operates.

To model the emergence of both spontaneous orientation map-like activity patterns as well as spontaneous OD map-like activity patterns, lateral connectivity between cortical sites was taken to be a sum of the contributions of ocular dominance sensitive synapses and orientation sensitive synapses, i.e.: 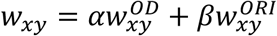

The contribution of orientation sensitive synapses to lateral connection was modeled following ^6^, in which connectivity was derived from the empirical polar map: If the animal is presented with several evenly spaced oriented gratings spanning the orientation domain (0-π), the information from all the experimental conditions can be used to derive the following map - 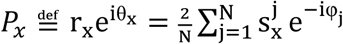

where θ_x_ is the preferred orientation of column at cortical location *x* and r_x_ its selectivity index. The map can then be used to determine the coupling between each pair of columns, i.e.: 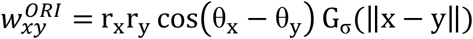

where 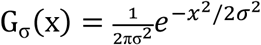, *σ* is given in mm and determines the spatial extent of the interactions. G_σ_ was set to 0 if ‖x − y‖ exceeded 2*σ* + 1in pixel units (each pixel was 120*μm*).

The contribution of ocular dominance synapses to lateral connections was determined by using the normalized empirical OD map according to:

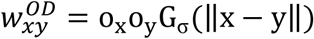

where o_x_ ∈ [−1,1], is the ocular preference.

For simplicity, spatial interactions were restricted spatially for both orientation synapses and OD synapses, to the same extent - i.e. using a single parameter *σ* - although some results suggest that this parameter may differ for both types of sensory information^47^.

The synaptic weights impinging on each column were normalized such that they would sum to 1, which is instrumental in maintaining the well-studied properties of each JR module within the context of an entire array of modules. Therefore, the weights *α, β* reflecting the relative contribution of orientation and OD synapses reflect the variance of weights for each type of information.

Similar to ^45^ spontaneous activity was induced by driving the network with iid equally distributed noise in the [0,300] interval.

## References

1 Arieli, A., Sterkin, A., Grinvald, A. & Aertsen, A. Dynamics of ongoing activity: explanation of the large variability in evoked cortical responses. Science 273, 1868–1871 (1996).

2 Tsodyks, M., Kenet, T., Grinvald, A. & Arieli, A. Linking spontaneous activity of single cortical neurons and the underlying functional architecture. Science 286, 1943–1946 (1999).

3 Kenet, T., Bibitchkov, D., Tsodyks, M., Grinvald, A. & Arieli, A. Spontaneously emerging cortical representations of visual attributes. Nature 425, 954–956 (2003).

4 O’Hashi, K. et al. Interhemispheric synchrony of spontaneous cortical states at the cortical column level. Cerebral Cortex, 1–14 (2017).

5 Omer, D. B., Fekete, T., Ulchin, Y., Hildesheim, R. & Grinvald, A. Dynamic Patterns of Spontaneous Ongoing Activity in the Visual Cortex of Anesthetized and Awake Monkeys are Different. Cerebral Cortex, bhy099–bhy099, doi:10.1093/cercor/bhy099 (2018).

6 Blumenfeld, B., Bibitchkov, D. & Tsodyks, M. Neural network model of the primary visual cortex: From functional architecture to lateral connectivity and back. Journal of computational neuroscience 20, 219–241 (2006).

7 Steriade, M., Nunez, A. & Amzica, F. A novel slow (< 1 Hz) oscillation of neocortical neurons in vivo: depolarizing and hyperpolarizing components. The Journal of Neuroscience 13, 3252–3265 (1993).

8 Steriade, M., Timofeev, I. & Grenier, F. Natural waking and sleep states: a view from inside neocortical neurons. Journal of neurophysiology 85, 1969–1985 (2001).

9 Ferrarelli, F. et al. Breakdown in cortical effective connectivity during midazolam-induced loss of consciousness. Proceedings of the National Academy of Sciences 107, 2681–2686 (2010).

10 Goldberg, J. A., Rokni, U. & Sompolinsky, H. Patterns of ongoing activity and the functional architecture of the primary visual cortex. Neuron 42, 489–500 (2004).

11 Fekete, T. et al. Critical dynamics, anesthesia and information integration: lessons from multi-scale criticality analysis of voltage imaging data. Neuroimage, doi:10.1016/j.neuroimage.2018.08.026 (2018).

12 Pal, D., Hambrecht-Wiedbusch, V., Silverstein, B. & Mashour, G. Electroencephalographic coherence and cortical acetylcholine during ketamine-induced unconsciousness. British journal of anaesthesia 114, 979–989 (2015).

13 Galán, R. F. On how network architecture determines the dominant patterns of spontaneous neural activity. PloS one 3, e2148 (2008).

14 Wagner, A. D. et al. Building Memories: Remembering and Forgetting of Verbal Experiences as Predicted by Brain Activity. Science 281, 1188–1191, doi:10.1126/science.281.5380.1188 (1998).

15 Pessoa, L., Gutierrez, E., Bandettini, P. A. & Ungerleider, L. G. Neural Correlates of Visual Working Memory: fMRI Amplitude Predicts Task Performance. Neuron 35, 975–987, doi:10.1016/S0896-6273(02)00817-6 (2002).

16 Ress, D. & Heeger, D. J. Neuronal correlates of perception in early visual cortex. Nature neuroscience 6, 414–420, doi:10.1038/nn1024 (2003).

17 Dehaene, S. & Changeux, J.-P. Ongoing Spontaneous Activity Controls Access to Consciousness: A Neuronal Model for Inattentional Blindness. PLoS Biol 3, e141, doi:10.1371/journal.pbio.0030141 (2005).

18 Pessoa, L. & Padmala, S. Quantitative prediction of perceptual decisions during near-threshold fear detection. Proceedings of the National Academy of Sciences of the United States of America 102, 5612–5617, doi:10.1073/pnas.0500566102 (2005).

19 Fox, M. D., Snyder, A. Z., Zacks, J. M. & Raichle, M. E. Coherent spontaneous activity accounts for trial-to-trial variability in human evoked brain responses. Nature Neuroscience 9, 23–25, doi:10.1038/nn1616 (2006).

20 Fox, M. D., Snyder, A. Z., Vincent, J. L. & Raichle, M. E. Intrinsic Fluctuations within Cortical Systems Account for Intertrial Variability in Human Behavior. Neuron 56, 171–184, doi:10.1016/j.neuron.2007.08.023 (2007).

21 Hesselmann, G., Kell, C. A., Eger, E. & Kleinschmidt, A. Spontaneous local variations in ongoing neural activity bias perceptual decisions. Proceedings of the National Academy of Sciences 105, 10984–10989, doi:10.1073/pnas.0712043105 (2008).

22 Hesselmann, G., Kell, C. a. & Kleinschmidt, A. Ongoing activity fluctuations in hMT+ bias the perception of coherent visual motion. The Journal of neuroscience: the official journal of the Society for Neuroscience 28, 14481–14485, doi:10.1523/JNEUROSCI.4398-08.2008 (2008).

23 Northoff, G., Qin, P. & Nakao, T. Rest-stimulus interaction in the brain: a review. Trends in Neurosciences 33, 277–284, doi:10.1016/j.tins.2010.02.006 (2010).

24 Raichle, M. E. Two views of brain function. Trends in Cognitive Sciences 14, 180–190, doi:10.1016/j.tics.2010.01.008 (2010).

25 Berkes, P., Orbán, G., Lengyel, M. & Fiser, J. Spontaneous Cortical Activity Reveals Hallmarks of an Optimal Internal Model of the Environment. Science 331, 83–87, doi:10.1126/science.1195870 (2011).

26 Civillico, E. F. & Contreras, D. Spatiotemporal properties of sensory responses in vivo are strongly dependent on network context. Frontiers in Systems Neuroscience 6, 25, doi:10.3389/fnsys.2012.00025 (2012).

27 Ponce-Alvarez, A., Thiele, A., Albright, T. D., Stoner, G. R. & Deco, G. Stimulus-dependent variability and noise correlations in cortical MT neurons. Proceedings of the National Academy of Sciences 110, 13162–13167, doi:10.1073/pnas.1300098110 (2013).

28 Singer, W. Cortical dynamics revisited. Trends Cogn Sci 17, 616–626, doi:10.1016/j.tics.2013.09.006 (2013).

29 Sachidhanandam, S., Sreenivasan, V., Kyriakatos, A., Kremer, Y. & Petersen, C. C. H. Membrane potential correlates of sensory perception in mouse barrel cortex. Nature Neuroscience 16, 1671–1677, doi:10.1038/nn.3532 (2013).

30 Zagha, E., Casale, Amanda E., Sachdev, Robert N. S., McGinley, Matthew J. & McCormick, David A. Motor Cortex Feedback Influences Sensory Processing by Modulating Network State. Neuron 79, 567–578, doi:10.1016/j.neuron.2013.06.008 (2013).

31 Zou, Q. et al. Intrinsic resting-state activity predicts working memory brain activation and behavioral performance. Human Brain Mapping 34, 3204–3215, doi:10.1002/hbm.22136 (2013).

32 Netser, S., Dutta, A. & Gutfreund, Y. Ongoing activity in the optic tectum is correlated on a trial-by-trial basis with the pupil dilation response. Journal of Neurophysiology 111, 918–929, doi:10.1152/jn.00527.2013 (2014).

33 Zhang, M., Wang, X. & Goldberg, M. E. A spatially nonselective baseline signal in parietal cortex reflects the probability of a monkey’s success on the current trial. Proceedings of the National Academy of Sciences 111, 8967–8972, doi:10.1073/pnas.1407540111 (2014).

34 Carcea, I., Insanally, M. N. & Froemke, R. C. Dynamics of auditory cortical activity during behavioural engagement and auditory perception. Nat Commun 8, 14412, doi:10.1038/ncomms14412 (2017).

35 Wilf, M. et al. Spontaneously Emerging Patterns in Human Visual Cortex Reflect Responses to Naturalistic Sensory Stimuli. Cereb Cortex 27, 750–763, doi:10.1093/cercor/bhv275 (2017).

36 Abbott, L. F. & Dayan, P. The Effect of Correlated Variability on the Accuracy of a Population Code. Neural Computation 11, 91–101, doi:10.1162/089976699300016827 (1999).

37 Arieli, A., Shoham, D., Hildesheim, R. & Grinvald, A. Coherent spatiotemporal patterns of ongoing activity revealed by real-time optical imaging coupled with single-unit recording in the cat visual cortex. Journal of Neurophysiology 73, 2072 –2093 (1995).

38 Gutnisky, D. A., Beaman, C. B., Lew, S. E. & Dragoi, V. Spontaneous Fluctuations in Visual Cortical Responses Influence Population Coding Accuracy. Cereb Cortex 27, 1409–1427, doi:10.1093/cercor/bhv312 (2017).

39 Omer, D. B., Hildesheim, R. & Grinvald, A. Temporally-structured acquisition of multidimensional optical imaging data facilitates visualization of elusive cortical representations in the behaving monkey. Neuroimage 82, 237–251, doi:10.1016/j.neuroimage.2013.05.045 (2013).

40 Arieli, A., Grinvald, A. & Slovin, H. Dural substitute for long-term imaging of cortical activity in behaving monkeys and its clinical implications. Journal of neuroscience methods 114, 119–133 (2002).

41 Shtoyerman, E., Arieli, A., Slovin, H., Vanzetta, I. & Grinvald, A. Long-term optical imaging and spectroscopy reveal mechanisms underlying the intrinsic signal and stability of cortical maps in V1 of behaving monkeys. The Journal of Neuroscience 20, 8111–8121 (2000).

42 Slovin, H., Arieli, A., Hildesheim, R. & Grinvald, A. Long-term voltage-sensitive dye imaging reveals cortical dynamics in behaving monkeys. Journal of neurophysiology 88, 3421–3438 (2002).

43 Fekete, T., Rubin, D., Carlson, J. M. & Mujica-Parodi, L. R. The NIRS Analysis Package: Noise Reduction and Statistical Inference. PloS one 6, e24322 (2011).

44 Calhoun, V., Adali, T., Pearlson, G. & Pekar, J. A method for making group inferences from functional MRI data using independent component analysis. Human brain mapping 14, 140–151 (2001).

45 Jansen, B. H. & Rit, V. G. Electroencephalogram and visual evoked potential generation in a mathematical model of coupled cortical columns. Biological cybernetics 73, 357–366 (1995).

46 Wendling, F., Bellanger, J.-J., Bartolomei, F. & Chauvel, P. Relevance of nonlinear lumped-parameter models in the analysis of depth-EEG epileptic signals. Biological cybernetics 83, 367–378 (2000).

47 Buzás, P. et al. Model-based analysis of excitatory lateral connections in the visual cortex. Journal of Comparative Neurology 499, 861–881 (2006).

